# Gating mass cytometry data by deep learning

**DOI:** 10.1101/054411

**Authors:** Huamin Li, Uri Shaham, Kelly P. Stanton, Yi Yao, Ruth Montgomery, Yuval Kluger

**Affiliations:** Applied Mathematics Program, Yale University, 51 Prospect St., New Haven, CT 06511, USA; Department of Statistics, Yale University, 24 Hillhouse Ave., New Haven, CT 06511, USA; Department of Pathology and Yale Cancer Center, Yale University School of Medicine, New Haven, CT, USA; Department of Internal Medicine, Yale School of Medicine, 333 Cedar St., New Haven, CT 06520, USA; Interdepartmental Program in Computational Biology and Bioinformatics,Yale University, New Haven, CT, USA

**Keywords:** FCM, CyTOF, automatic gating, deep learning, domain adaptation

## Abstract

Mass cytometry or CyTOF is an emerging technology for high-dimensional multiparameter single cell analysis that overcomes many limitations of fluorescence-based flow cytometry. New methods for analyzing CyTOF data attempt to improve automation, scalability, performance, and interpretation of data generated in large studies. Assigning individual cells into discrete groups of cell types (gating) involves time-consuming sequential manual steps, untenable for larger studies. We introduce DeepCyTOF, a standardization approach for gating, based on deep learning techniques. DeepCyTOF requires labeled cells from only a single sample. It is based on domain adaptation principles and is a generalization of previous work that allows us to calibrate between a target distribution and a source distribution in an unsupervised manner. We show that Deep-CyTOF is highly concordant (98%) with cell classification obtained by individual manual gating of each sample when applied to a collection of 16 biological replicates of primary immune blood cells, even when measured accross several instruments. Further, DeepCyTOF achieves very high accuracy on the semi-automated gating challenge of the FlowCAP-I competition as well as two CyTOF datasets generated from primary immune blood cells: (i)14 subjects with a history of infection with West Nile virus (WNV), (ii) 34 healthy subjects of different ages. We conclude that deep learning in general, and DeepCyTOF specifically, offers a powerful computational approach for semi-automated gating of CyTOF and flow cytometry data.

## 1 Introduction

Flow cytometry (FCM) is routinely used in cellular and clinical immunology. Current fluorescence-based FCM experiments provide up to 15 numeric parameters for each individual cell from blood samples in a high-throughput fashion. This allows efficient multiparameter characterization of single cell states. Interpretation of such data from hundreds-of-thousands to millions of cells is paramount to understanding the pathogenesis of a broad range of human diseases. Mass cytometry (CyTOF) is an emergent technological development for high-dimensional multiparameter single cell analysis. By using heavy metal ions as labels and mass spectrometry as readout, many more markers can be simultaneously measured. Current CyTOF instruments allow users to probe over 40 antibody specificities and thus provide a significant improvement in analyzing the underlying cell sub-populations of a system [1, 2]. CyTOF provides unprecedented multidimensional immune cell profiling and has recently been applied to characterizing peripheral blood cells, Natural Killer cells in viral infections, skin cells, cells in celiac disease, responding phenotypes in cancer, and even holds the promise of examination of solid tumors [3, 4, 5, 6, 7, 8, 9, 10]. Cellular characterization by FCM and CyTOF will improve our understanding of disease processes [11].

Gating (assigning individual cells into discrete groups of cell types) is one of the important steps and a bottleneck of analyzing FCM and CyTOF data. It involves time-consuming sequential manual steps, untenable for larger studies [12, 13, 14, 15, 16, 17, 18, 19]. The time it takes to manually analyze a cytometry experiment depends on the number of blood samples as well as the number of markers [20]. Specifically, gating is performed by drawing polygons in the plane of every two markers. This implies that the time required for gating is roughly quadratic in the number of markers. In addition, the manual procedure, combined with the increase in the number of markers, make this process prone to human errors. Technical variation naturally arises due to the variation between individual operators [21]. The subjectivity of manual gating introduces variability into the data and impacts reproducibility and comparability of results, particularly in multi-center studies [22]. Thus the slow processing time and the inherent subjectivity of manual analysis should be considered as primary reasons for using computational assistance methods.

The FlowCAP consortium aims to boost user confidence in the viability of automated gating methods [23]. Many of the pipelines described therein are tailored for exploratory, discovery-oriented data analysis. New methods for analyzing cytometry data continue to emerge; these methods attempt to improve automation, scalability, performance, and interpretation of data generated in large studies. These computational methods can be categorized as unsupervised or supervised approaches. Both types of approaches use a variety of simple linear transformations, density estimations, hierarchical clustering, and nonlinear projection methods, that together allow extracting features that can be used to study differences between samples^1^. However, most current tools are less suitable for routine use where analysis must be standardized, reproducible, interpretable, and comparable [24]. In general, no automated gating algorithm or approach that would solve all specific computational problems has been accepted as the gold standard for replacing manual gating [23, 25].

Additionally, in the last few years, deep learning methods have achieved outstanding performance in various computational tasks, such as image analysis, natural language processing, and pattern recognition [26]. These approaches have also been shown to be effective for extracting natural features from data in general settings [27, 28, 29]. Moreover, recent efforts to use deep learning approaches in genomics and biomedical applications show their flexibility for handling complex problems [30, 31, 32, 33, 34]. However, deep learning typically requires very large numbers of training instances and thus its utility for many genomic, proteomic and other biological applications is questionable. While in most genomics applications, the number of instances (e.g., number of gene expression arrays) is typically smaller than the number of variables (e.g., genes), in each FCM and CyTOF run we typically collect approximately 10^5^ to 10^6^ cells, so that the number of instances (cells) is several orders of magnitude larger than the number of variables (up to 50 markers). Therefore, developing deep learning approaches to analyze cytometry data is very promising.

However, in FCM and CyTOF experiments, variation in both biological and technical sources can make automatic gating challenging. Instrument calibration causes variation across samples, such a situation is often referred to “batch effect”. In order to avoid gating each dataset separately (which therefore requires labeled samples from each dataset), a domain adaptation procedure is used. *Domain Adaptation* is a set of techniques that allow the use of a learning scheme (or model) trained on data from a source domain with a given distribution, which can then be applied to a target domain with a related but not equivalent distribution. The objective of domain adaptation is to minimize the generalization error of instances from the target domain [35, 36].

We present DeepCyTOF, an integrated deep learning domain adaptation framework, which employs one manually gated reference sample and utilizes it for automated gating of the remaining samples in a study. We first include two preprocessing options to use a denoising autoencoder (DAE) to handle missing data and use multiple distribution-matching residual networks (MMD-ResNets) [37] to calibrate an arbitrary number of source samples to a fixed reference sample, and then perform a domain adaptation procedure for automatic gating.

We demonstrate DeepCyTOF for automatic gating of three CyTOF datasets consisting of 56, 136 and 16 samples respectively, and obtain almost identical results to those obtained by manual gating. In particular, the collection of 16 samples were obtained from the same subject and measured on multiple CyTOF instruments, a setting where batch effects are strong. We demonstrate that by using DeepCyTOF’s preprocessing options to account for the missing data and the batch effects, one is able to utilize manual gating of a single source sample to perform high quality automatic gating of the remaining samples, and significantly reduce the time and effort that are currently required for manual gating. In addition, we compare DeepCyTOF to the other competing supervised approaches benchmarked on each dataset of the 4th challenge of the FlowCAP-I competition [23], and show that DeepCyTOF outperforms each dataset’s respective winning algorithm, achieving state-of-the-art accuracy.

The structure of this manuscript is as follows: in Section 2 we describe the datasets and algorithms used in this research. Experimental results are given in Section 3. In Section 4 we explain the theoretical justification of Deep-CyTOF as a domain adaptation procedure. Section 5 concludes this manuscript.

## 2 Materials and Methods

### 2.1 Datasets

#### 2.1.1 FlowCAP-I datasets

We employ five collections of FCM datasets from FlowCAP-I [23]: (1) Diffuse large B-cell lymphoma (DLBCL), (2) Symptomatic West Nile virus (WNV), (3) Normal donors (ND), (4) Hematopoietic stem cell transplant (HSCT), and (5) Graft-versus-host disease (GvHD). With the results from manual gates produced by expert analysis, the goal of FlowCAP-I challenges is to compare the results of assigning cell events to discrete cell populations using automated gates. In particular, we consider Challenge 4: supervised approaches trained using human-provided gates. We use the manual gating provided from FlowCAP-I to evaluate the neural nets predictions.

#### 2.1.2 Mass Cytometry Datasets

We analyze two collections of CyTOF datasets measured on one instrument in the Montgomery Lab. The datasets consist of primary immune cells from blood of (1) *N* = 14 subjects (8 asymptomatic and 6 severe) with a history of infection with West Nile virus (WNV), and (2) *N* = 34 healthy subjects of different ages (20 young and 14 old). Each blood sample is labeled with *d* = 42 antibody markers [5], 12 of which are used in our analysis as they are the relevant markers for the task of classification described below: HLA-DR, CD4, CD8, CD3-UCH1, CD16, CD33, CD19, CD14, CD56, DNA1, DNA2, Cisplatin. Each sample is subjected to four CyTOF experiments including a baseline state and three different stimuli (PMA/ionomycin, tumor cell line K562, and infection with WNV). The goal is to classify each cell to one of six cell type categories: (1) B cell, (2) CD4+ T cell, (3) CD8+ T cell, (4) Monocytes, (5) Natural killer (NK) cells, and (6) unlabeled. There are 56 and 136 samples in the first two datasets, and we manually gate each single cell to evaluate the neural nets predictions.

We analyze an additional third collection of 16 CyTOF samples, which are all drawn from a single subject. These samples are measured in two different times and instruments as a part of a multi-center study [38]. The first 8 samples are collected at the same time, and the last 8 samples are collected two months apart from the first 8. Additionally, each consecutive two samples are measured by the same instrument. Each of the 16 samples contains 26 markers, out of which eight correspond to the following protein markers: CCR6, CD20, CD45, CD14, CD16, CD8, CD3, CD4; we perform our classification experiments on this 8-dimensional data as these eight markers are the relevant ones for the task of classification: classify each cell to one of five cell type categories: (1) B cell, (2) CD4+ T cell, (3) CD8+ T cell, (4) Monocytes, and (5) unlabeled. We manually gate each single cell to evaluate the neural nets predictions.

#### 2.1.3 Pre-processing

For FlowCAP-I datasets, we apply a logarithmic transform, followed by rescaling, as described in Appendix A. For Mass Cytometry datasets, we first manually filter all samples to remove debris and dead cells. In addition, different samples are measured at different times; fine changes in the state of the CyTOF instrument between these runs introduce additional variability into the measurements (batch effects). The specific nature of these changes is neither known nor modeled. To tackle this problem and apply a gating procedure, we follow most practitioners in the field, and calibrate the samples by applying an experimental-based normalization procedure^2^. This procedure involves mixing samples with polystyrene beads embedded with metal lanthanides, followed by an algorithm which enables correction of both short-term and long-term signal fluctuations [39]. Once the data is normalized, we apply a logarithmic transform, and rescaling.

### 2.2 DeepCyTOF Algorithm

DeepCyTOF integrates between three different tasks needed to achieve automated gating of cells in multiple target samples^3^ based on manual gating of a single reference source sample. The tasks include sample denoising, calibration between target samples and a single reference source sample and finally cell classification. We implement each of these tasks using the following three neural nets: (1) a denoising autoencoder (DAE) for handling missing data; (2) an MMD-ResNet for calibrating between the target samples and a reference source sample; (3) a depth-4 feed-forward neural net for classifying/gating cell types trained on a reference source sample. DeeopCyTOF has options to run with or without denoising, and with or without calibration. In examples shown in the Results section DeepCyTOF was employed to classify cells without denoising and without calibration (Section 3.1), without calibration but with the denoising preprocessing step (Section 3.2) and with the denoising and with the calibration preprocessing steps (Section 3.3).

#### 2.2.1 Removing Zeros using Denoising Autoencoder

All samples in our Mass Cytometry dataset contain large proportions of zero values across different markers. This usually does not reflect biological phenomenon, but rather, occurs due to instabilities of the CyTOF instrument. To tackle this, we include an option in DeepCyTOF to remove the zeros by training a denoising autoencoder (DAE) [40] on the cells with no or very few zero values. A DAE is a small neural net that is trained to reconstruct a clean input from its corrupted version. Unlike [40], who use Gaussian noise to corrupt the inputs, we use dropout noise, i.e., we randomly zero out subset of the entries of each cell, to simulate the machine instabilities. We train a DAE for each batch, by combining all samples from that batch, selecting the cells with no zeros and using them as training set. For each DAE, we set the dropout probability to be the proportion of zeros in the measurement of the corresponding batch. Once a DAE is trained, we pass all samples from its batch through it to denoise the data.

#### 2.2.2 MMD-ResNet

To account for machine based technical bias and variability, we include a preprocessing option in DeepCyTOF to calibrate each batch to a reference using the MMD-ResNet approach. MMD-ResNet [37] is a deep learning approach to learn a map that calibrates the distribution of a source sample to match that of a target sample. It is based on a residual net (ResNets) architecture [41, 42] and has Maximum Mean discrepancy (MMD) [43, 44] as the loss function. ResNet is a highly successful deep networks architecture, which is based on the ability to learn functions which are close to the identity. MMD is a measure for a distance between distributions, which had been shown to be suitable for training of neural nets-based generative models [45, 46]. If 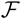 is a reproducing kernel Hilbert space with a (universal) kernel function k(·, ·), the (squared) MMD between distributions *p, q* over a space 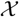 is defined as

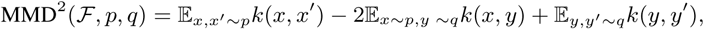

where *x* and *x′* are independent, and so are *y* and *y′*.

For calibration purposes, we want to find a map that brings the distribution of the source sample close to that of the target sample; we further assume that this map should be close to the identity. In a previous work [37], we have shown that MMD-ResNets are successful in learning such maps, and used them for calibration of CyTOF data and single cell RNA sequencing data. We refer the reader to [37] for a more comprehensive description of MMD-ResNets.

In this manuscript, we follow the approach of [37] and use ResNets consisting of three blocks, which are trained to minimizing the loss

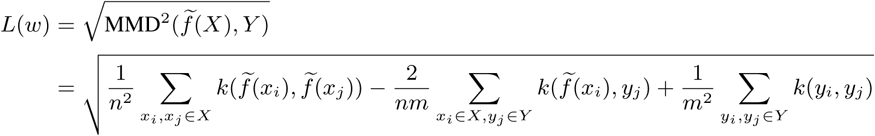

where 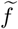 is the map computed by the network, w are the network parameters, and 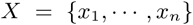, 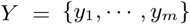 are two finite samples from the source and target distributions, respectively. An example of a MMD-ResNet is shown in Figure 2.

#### 2.2.3 Cell Classifier

To choose which sample will be used as reference source sample, for each sample i we first compute the *d* × *d* covariance matrix Σ*_i_*, where *d* is the number of markers (dimensionality) of these samples. For every two samples *i*, *j* we then compute the Frobenius norm of the difference between their covariance matrices ║Σ*_i_* − Σ*_j_*║_*F*_, and we select the sample with the smallest average distance to all other samples to be the reference sample. Once the reference sample is chosen, we use manual gating to label its cells; the gating is used as ground truth labels to train the classifier. This is the only label information DeepCyTOF requires, regardless of the total number of samples we want to gate.

To classify cell types, we trained depth-4 feed-forward neural nets, each consisting of three softplus hidden layers and a softmax output layer. Further technical details regarding the architecture and training are given in Section 3.4. An example of such classifier is shown in Figure 1.

**Figure 1:**
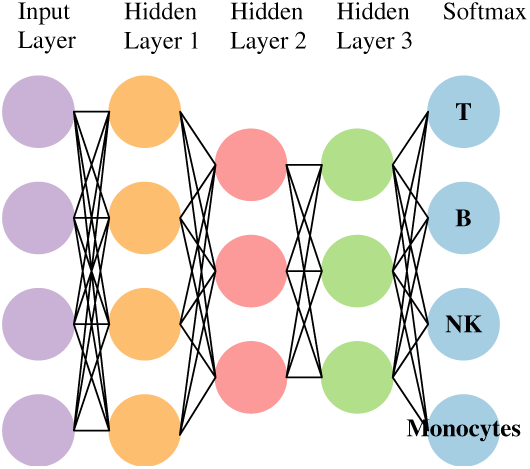
A neural net for classifying cell types.

**Figure 2:**
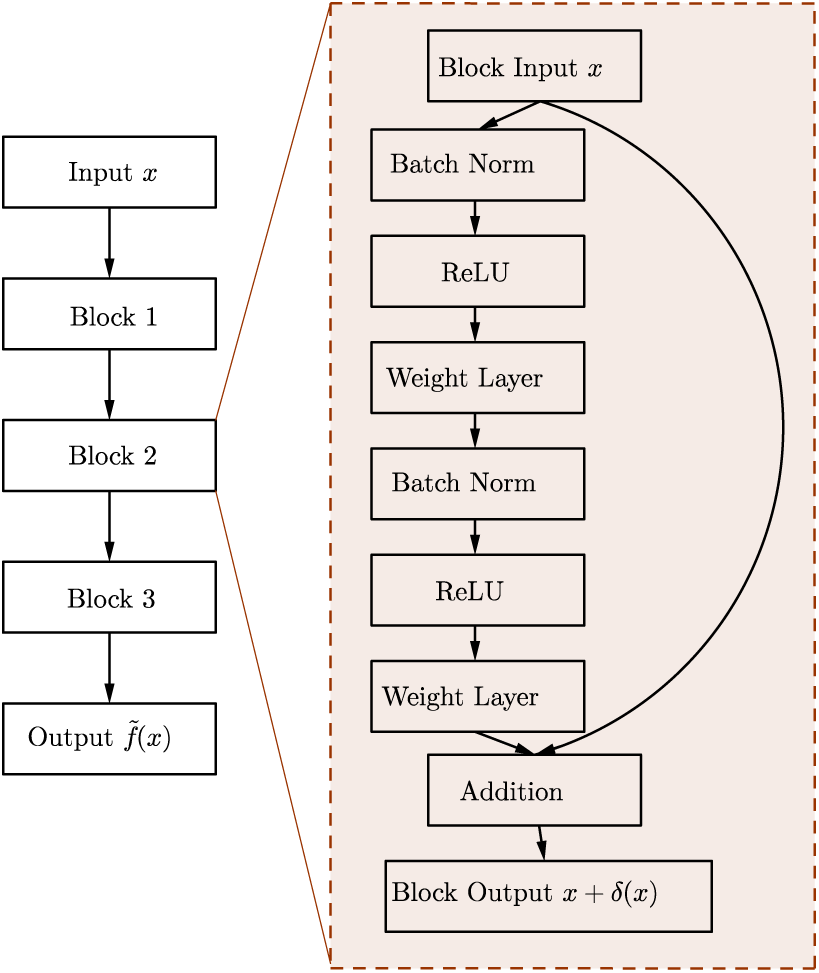
MMD-ResNet with 3 blocks.

### 2.3 Evaluation

To compare DeepCyTOF approach to algorithms from the 4th challenge of the FlowCAP-I competition, 25% of the cells of each subject from the FCM datasets in FlowCAP-I are labeled by manual gating and used to train a cell type classifier based on the procedure of Section 2.2.3, which is then used to predict the labels of the remaining 75% cells. Here the DAE option is disabled because there are no missing values and the MMD-ResNet option is also disabled because the training and test sets are from the same run and thus do not require calibration.

To perform semi-automated gating of all samples of each of the first two CyTOF datasets based on the procedure of Section 2.2, we select a single baseline reference sample as in Section 2.1.2. We manually gate this sample, and use it to train a classifier for predicting the cell type class of each cell. The other baseline samples and additional samples that undergo three different stimuli (PMA/ionomycin, tumor cell line K562, and infection with WNV) are left for testing. Batch effects in these two datasets were not substantial allowing classification of cells into major cell populations without employing a calibration step as in Section 2.2.2 prior for to the cell classification of the test samples.

To perform semi-automated gating of the samples in the third multi-center CyTOF dataset, we follow the procedure of Section 2.2, i.e., we choose a single reference sample, train a collection of MMD-ResNets to calibrate all the remaining samples to it, train a cell type classifier on manually gated data of the reference sample, and use it to classify the cells of the calibrated samples. We compare the classification performance between two options of running DeepCyTOF. One option is with calibration of the target samples by MMD-ResNets and the other option is without calibration.

We use the F-measure statistic (the harmonic mean of precision and recall) for the evaluation of our methods as described in [23]. The F-measure for multiple classes is defined as the weighted average of F-measures for each cell type, i.e.,

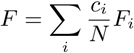

where *c_i_* is the number of cells with type *i*, *N* is the total number of cells, *F_i_* is the F-measure for the *i*th cell type versus all other types (including unknown types):

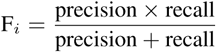

An F-measure of 1.0 indicates perfect agreement with the labels obtained by manual gating. For any given vector of F-measure values on a given dataset, we create several vectors of F-measure values (by sampling with replacement), compute the mean F-measure and 95% bootstrap percentile confidence interval for the mean.

## 3 Results

In this section, we present results from three experiments: (1) cell classification of five FCM datasets from the FlowCAP-I competition by applying DeepCyTOF without denoising (DAE) and without calibration (MMD-ResNets), as they are not needed as explained above, (2) cell classfication of two CyTOF datasets by applying DeepCyTOF with the denoising option (DAE), but without calibration (MMD-ResNets) of the target samples, (3) cell classfication of a multi-center CyTOF dataset by applying DeepCyTOF with the denoising option (DAE), and with calibration (MMD-ResNets) of the target samples to the source sample.

### 3.1 Evaluation of classification performance from FlowCAP-I

Table 1 presents the performance of DeepCyTOF when applied to the five datasets from the 4th challenge of FlowCAP-I competition. The performance is compared to the respective winner of each of the five collections in this competition. As can be seen, our predictions are better than the competition winner in four out of the five collections and similar on the HSCT collection.

**Table 1:** Summary of results for the FlowCAP-I cell identification challenge. The numbers in parentheses represent 95% confidence intervals

| $F = \frac{\text{precision} \times \text{recall}}{\text{precision} + \text{recall}}$ | DeepCyTOF<sup>a</sup> | Competition's winner |
| --- | --- | --- |
| GvHD<sup>b</sup> | 0.986 (0.979, 0.991) | 0.92 (0.88, 0.95) |
| DLBCL | 0.985 (0.976, 0.993) | 0.95 (0.93, 0.97) |
| HSCT | 0.991 (0.988, 0.993) | 0.98 (0.96, 0.99) |
| WNV | 0.999 (0.998, 0.999) | 0.96 (0.94, 0.97) |
| ND | 0.988 (0.987, 0.989) | 0.94 (0.92, 0.95) |

<sup>a</sup> DeepCyTOF without denoising (DAE) and without calibration (MMD-ResNets).
<sup>b</sup> FlowCAP-I collections: graft-versus-hist disease (GvHD); diffuse large B-cell lymphoma (DLBCL); symptomatic West Nile virus (WNV); normal donors (ND); hematopoietic stem cell transplant (HSCT).

### 3.2 Application of DeepCyTOF to CyTOF datasets in the Absence of Strong Batch Effects

In this experiment, for each of the two different collections (which contain 14 and 34 baseline samples, respectively), we chose a reference sample, used it to train DeepCyTOF using the options that omits the calibration step, which was then used to predict the cell types in all the other samples in that collection (55 samples from the Asymptomatic vs. Severe WNV dataset and 135 samples from the Old vs. Young dataset), without any calibration.

Figure 3 shows an example of embedding of labeled cells in a three dimensional space, obtained from the top hidden layer of a neural net cell type classifier, as the ones used for this experiment. As can be seen, most of the labeled cells concentrate in separated clusters representing specific cell types. Table 2 summarizes the results, and provides a comparison to a shallow, linear classifier (softmax). Table 2 illustrates some interesting points: first, nearly perfect performance on the test data is achieved, despite the fact that it was only given labels from the reference sample. Second, DeepCyTOF performs significantly better than softmax regression, which may be a result of the depth and the non-linearity of the network. Third, whether or not the data is normalized does not affect the performance of DeepCyTOF. Fourth, we also applied DeepCyTOF choosing the option that includes a preprocessing calibration step in which MMD-ResNets are used to calibrate the source samples to the reference sample. However, this yields a modest improvement in the results. This may be due to the fact that the datasets did not significantly differ in distribution. A different scenario, where significant differences exist in the data, is considered in Section 3.3.

**Figure 3:**
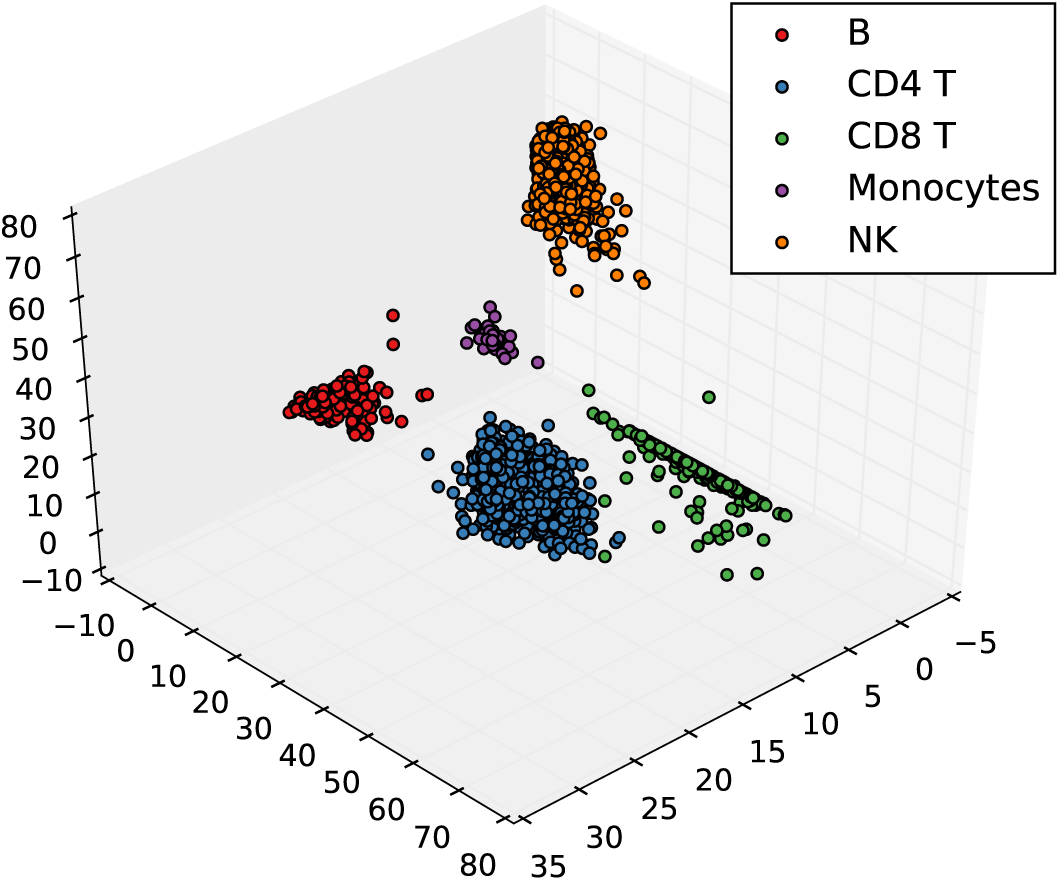
Third hidden layer representation of a blood sample (the unlabeled cells are omitted), obtained from DeepCyTOF without a calibraition step. Each color corresponds to a cell type. Different cell types are concentrated in different regions of the code space.

**Table 2:** Summary of results for the two CyTOF collections. The numbers in parentheses represent 95% confidence intervals.

| $F = \frac{\text{precision} \times \text{recall}}{\text{precision} + \text{recall}}$ | AnS-UN<sup>a</sup> | AnS-N | OnY-UN | OnY-N |
| --- | --- | --- | --- | --- |
| DeepCyTOF<sup>b</sup> | 0.990 (0.987, 0.993) | 0.991 (0.987, 0.994) | 0.993 (0.990, 0.997) | 0.993 (0.989, 0.996) |
| Softmax regression | 0.966 (0.955, 0.975) | 0.967 (0.957, 0.976) | 0.963 (0.946, 0.977) | 0.963 (0.946, 0.977) |

<sup>a</sup> Datasets: unnormalized Asymptomatic&Severe (AnS-UN); normalized Asymptomatic&Severe (AnS-N); normalized Old&Young (OnY-N); unnormalized Old&Young (OnY-UN).
<sup>b</sup> DeepCyTOF without using MMD-ResNets for calibration of the target samples.

### 3.3 Overcoming Strong Batch Effects using DeepCyTOF

The multi-center dataset consists of samples of the same subject, which were measured on different CyTOF machines in different locations. It is therefore reasonable that the different instrument conditions will result in calibration differences, which need to be accounted for. A domain adaptation framework like DeepCyTOF might be valuable in such scenario.

As in section 2.2, we first chose a reference subject (sample 2) as the source. For each target sample, we then trained a MMD-ResNet to calibrate it to the reference sample. Subsequently, we trained a cell classifier using labeled data from the reference sample, and used it to classify cells from all the remaining calibrated 15 samples. We compared the performance of DeepCyTOF to a similar procedure where we skip the calibration step so that the input to the cell classifier is the data from the un-calibrated target samples. Figure 4 shows the F-measure scores for each sample before and after calibration. As can be seen, the F-measure scores of samples 9-16 are significantly higher when a calibration step is included in the gating process. For samples 1-8 we observed that the scores are very high even without a calibration step. Overall, applying DeepCyTOF with a calibration step results in an weighted average F-measure of 0.985 with 95% confidence interval (0.979, 0.990), which is significantly higher compared to the weighted average F-measure obtained by applying DeepCyTOF without calibration, which was 0.925 with 95% confidence interval (0.890, 0.956).

**Figure 4:**
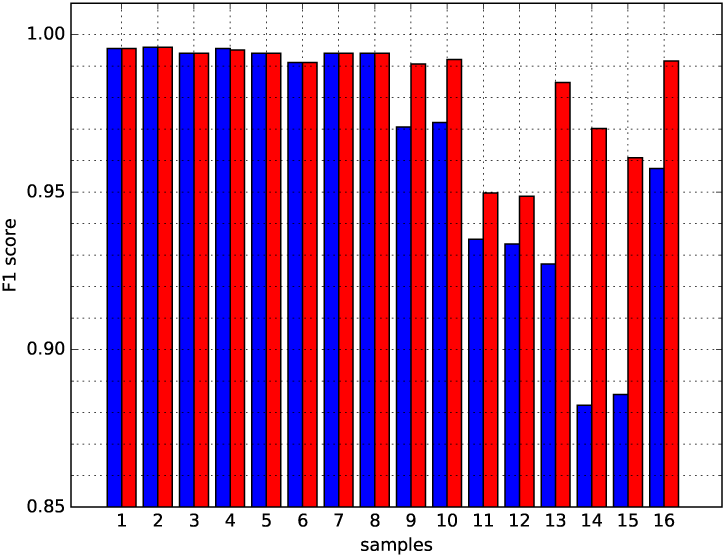
F-measure scores of 16 samples in the multi-center dataset before (blue) and after (red) calibration.

**Figure 5:**
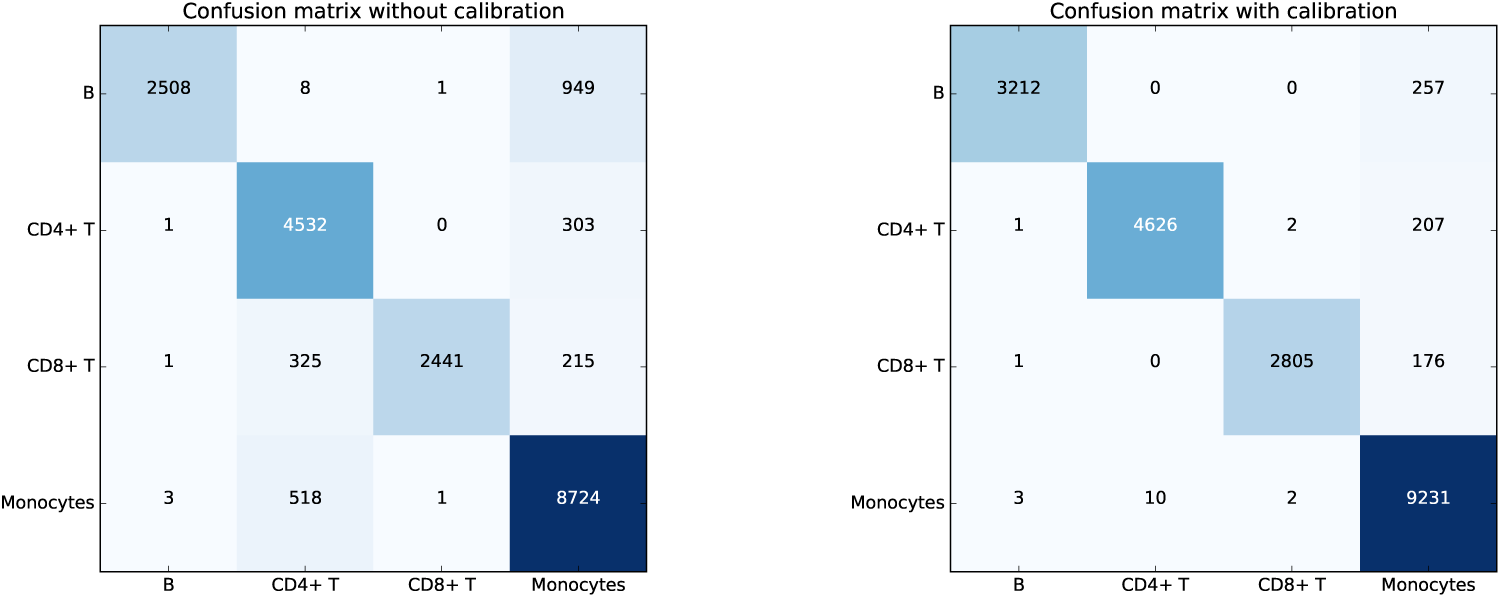
Confusion matrix for visualizing the performance of DeepCyTOF when applied to sample 15 without (left) and with (right) a calibration step (the unlabeled cells are omitted). The rows represent the actual cell type label and the columns represent the predicted cell type label. The F measure associated with the left panel (0.8857) is significantly lower than the F measure associated with the right panel (0.9609).

Figure 6 shows *t*-SNE embedding [47] of the cells of a representative sample selected from samples 1-8 (sample 7) and a representative sample from the other eight samples (sample 15) versus the cells of the reference sample, before and after calibration. On both samples the MMD-ResNet calibration seem to correct the batch effect appropriately, as after the calibration same cell types are embedded in the same clusters. To understand why the calibration almost did not change the accuracy on the last eight samples (while improving it dramatically on the last eight), Figure 7 in the supplementary material shows the MMD between the source sample and each of the target samples^4^. As we can see, in all samples (with the exception of sample 2, which is the source sample, and sample 1 which was measured on the same instrument as sample 2), the MMD-ResNet calibration reduces the MMD between the distributions. However, before calibration the MMD between each the first eight target samples and the reference sample is relatively small, possibly making the classifier generalize well to these distributions even without calibration.

**Figure 6:**
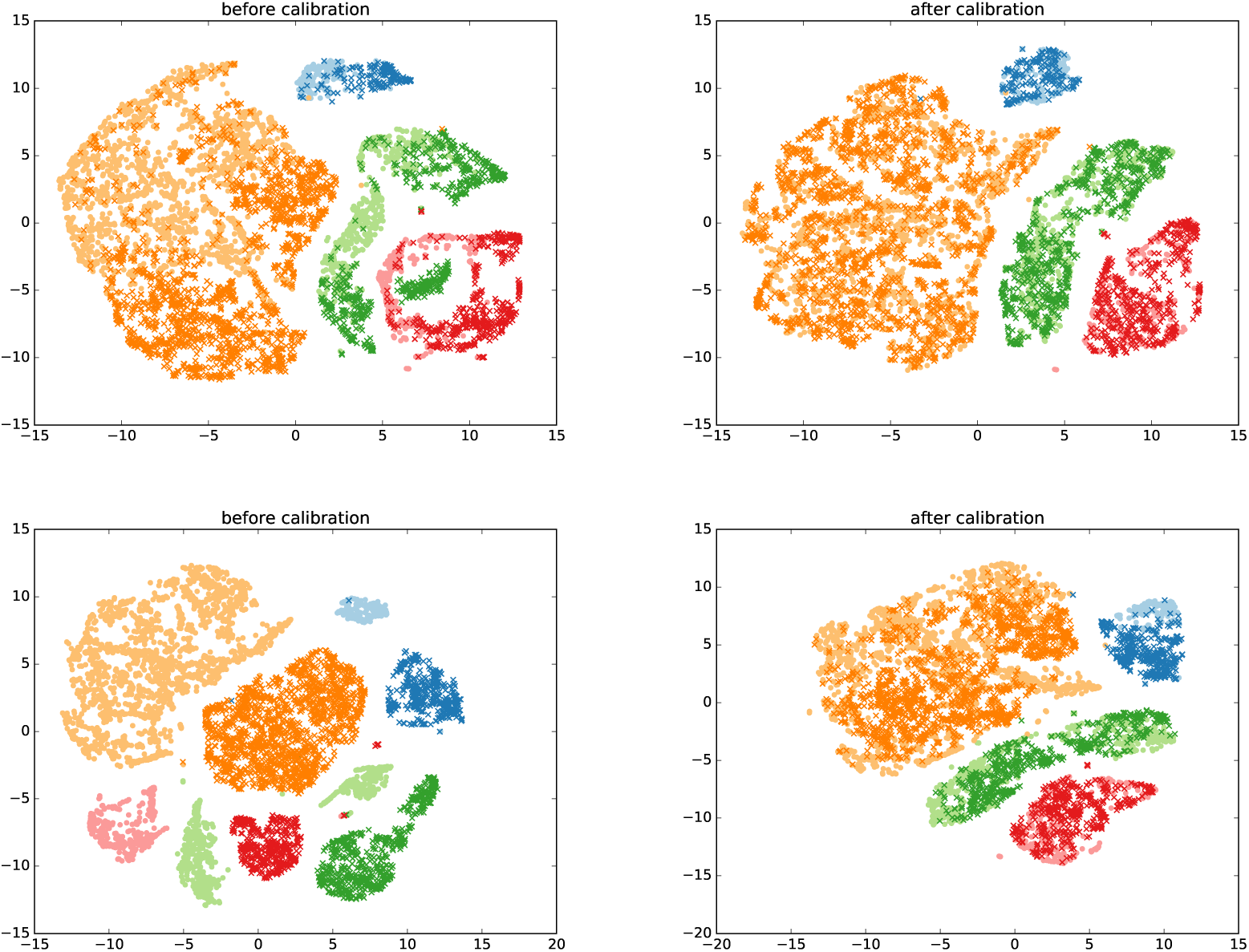
Top: t-SNE plots of the joint distribution of sample 7 (dark crosses) and the target sample (light circles) before (left) and after (right) calibration (the unlabeled cells are omitted). Bottom: Similarly to the upper panel but plots correspond to to the joint distribution of sample 15 and the target sample. Different cell types have different colors: B cells (light and dark blue), CD4+ T cells (light and dark green), CD8+ T cells (light and dark red), Monocytes cells (light and dark orange). After calibration, same cell types are clustered together.

**Figure 7:**
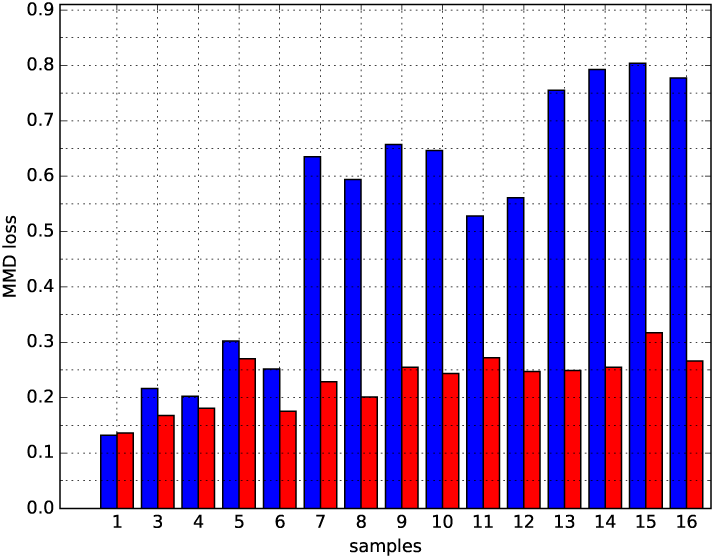
MMD between the source (sample 2) and each of the of other 15 samples in the multi-center dataset before (blue) and after (red) calibration. The MMD values were computed based on random batches of size 1000 from each sample.

In the supplementary material, we also provide additional results from this experiment. A more detailed perspective on the effect of the calibration on the classification accuracy for the samples 9-16 is given in Figure 5, which shows the confusion matrix of a representative sample (sample 15), obtained before and after the calibration. For this sample, the F-measure obtained by applying DeepCyTOF with and without a calibration step are, 0.9614 and 0.6603 respectively. In order to demonstrate the quality of calibration not only in a macroscopic level, but also when restricting the attention to a specific cell population, Figure 8 in the appendix shows the *t*-SNE plot of CD8+T cells from sample 15 before and after calibration. The results are consistent with the ones given above.

**Figure 8:**
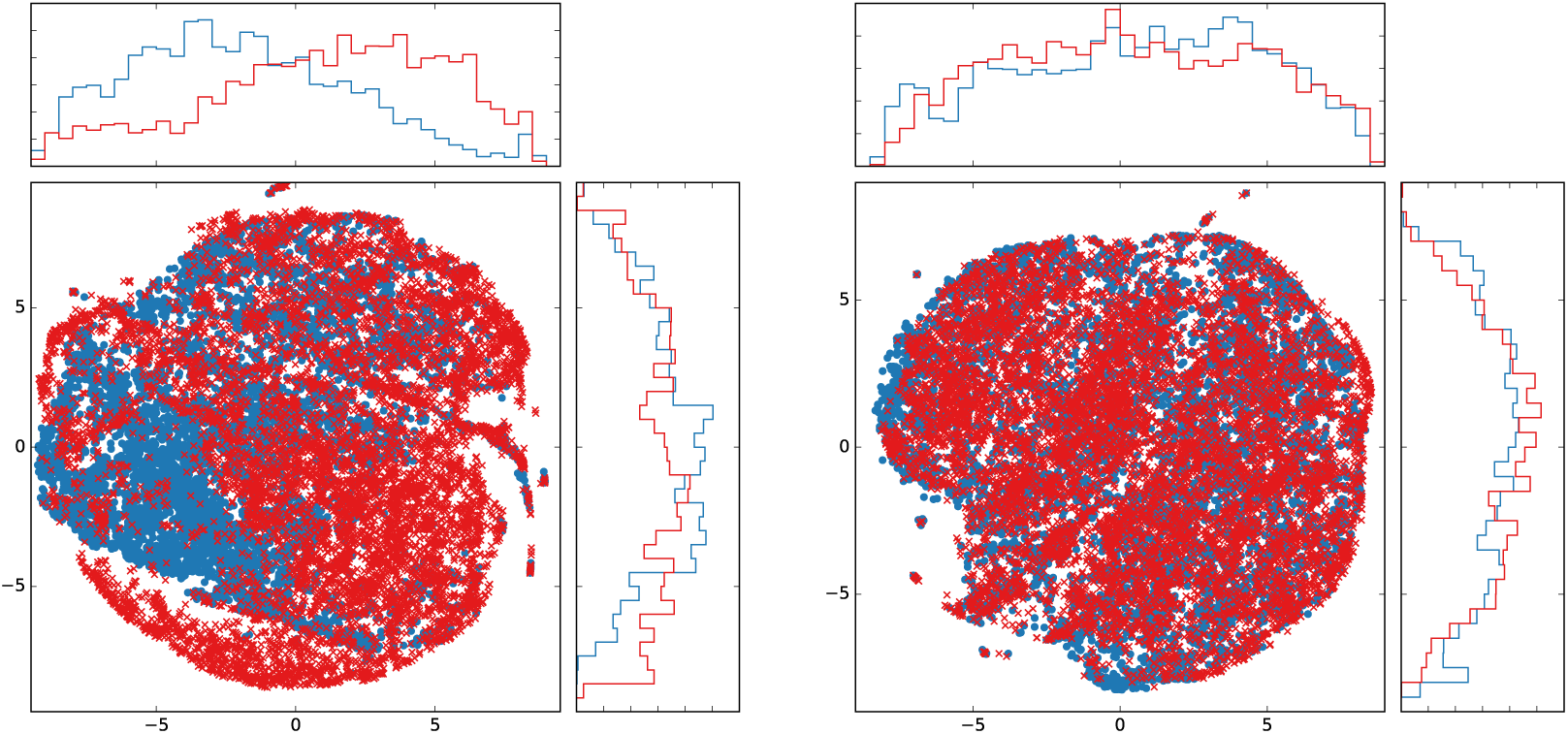
t-SNE plot of CD8+ T cell from sample 15 before (top) and after (bottom) with DeepCyTOF.

Finally, to demonstrate the effect of denoising on the quality of calibration, Figure **??** shows the MMD between the reference sample and each of the other samples with and without denoising. We see that with denoising the MMD is smaller than without denoising.

### 3.4 Technical Details

All cell type classifiers used in this work, were depth 4 feed-forward nets, with softplus hidden units and a softmax output layer, where the hidden layer sizes have been set to 12, 6, and 3. All MMD-ResNets were identical and consisted of three blocks, as can be seen in Figure 2. The first weight matrix was of size 25 × 8, and the second weight matrix was of size 8×25. The DAE hidden layer consisted of 25 ReLU units. All networks were trained using RMSprop [48]. We use the default learning rate (10^-2^) to train the DAE. The learning rate schedule for training the classifiers and MMD-ResNets was defined as follows:

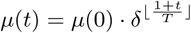

where *μ*(*t*) was the learning rate at epoch *t*, *μ*(0) was the initial learning rate, *δ* was a constant, and *T* was the schedule. For training the classifiers, we have *μ*(0) = 10^−3^, *δ* = .5, and *T* = 50. For training the MMD-ResNets, we have *μ*(0) = 10^−3^, *δ* = .1, and *T* = 15.

We used mini-batches of size 128 for the cell type classifiers and the DAE, and 1000 for the MMD-ResNets. For each net, a subset 10% of the training data is held out for validation, to determine when to stop the training. In the DAE and cell type classifiers, a penalty of 10^−4^ on the *l*_2_ norm of the network weights is added to the loss for regularization. In the MMD-ResNets we used for this purpose a penalty of 10^−2^.

The kernel used for MMD-ResNets is a sum of three Gaussian kernels

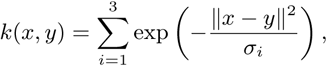

We set the scales for the Gaussian kernels of the MMD-ResNets to be *σ_i_*s to be 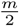, *m*, 2*m*, where m is the median of the average distance between a point in the target sample to its 25 nearest neighbors.

DeepCyTOF, implemented in Keras, is publicly available at https://github.com/hl475/DeepCyTOF.git.

## 4 Theoretical Justification for the Calibration Step of DeepCyTOF

In the classical domain adaptation setting [49], a domain is a pair 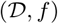, where 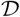 is a distribution on an input space 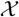 and 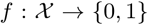 is a labeling function. A hypothesis is a function 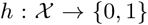. Given a domain 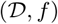, any hypothesis is associated with an error

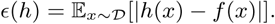

Given a hypothesis, a source domain 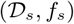 and a target domain 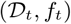, one is interested in expressing the target error

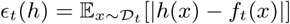

in terms of the source error

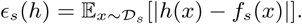

This corresponds to expressing the error of a classifier, trained on the source data, on the target data.

Let 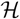 be hypothesis class of finite VC dimension. Ben David et al. [49] prove that for every 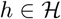, with a probability of at least 1 − *δ*,

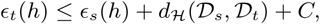

where

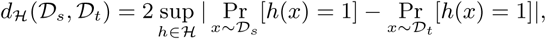

and *C* is a constant which does not depend on *h*.

By considering 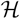 to be the class of Parzen window classifiers, it can be shown [50, 51] that

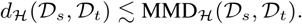

This implies that calibrating the data by minimizing the MMD between the source and target distribution, the target error will get close to the source error. This is precisely the procedure DeepCyTOF performs.

## 5 Discussion

In this work, we show that deep learning machinery can be very effective in classification of cell types; the performance substantially surpasses the predictive accuracy of the methods presented in the 4th challenge of the FlowCAP-I competition. In addition, we introduce DeepCyTOF, an automated framework for gating cell populations in cytometry samples. DeepCyTOF integrates deep learning and domain-adaption concepts. The labels obtained by manual gating of the reference sample are utilized in a domain-adaptation manner. These steps enable DeepCyTOF to inherently calibrate the major cell populations of multiple samples with respect to the corresponding cell populations of the reference sample. We analyze 208 CyTOF samples and observed nearly identical results to those obtained by manual gating (with mean F-measure ≥ 0.98).

In practice, run-to-run variations in CyTOF experiments both in the same instrument and between instruments are very common. These variations lead to significant batch effects in the datasets with samples collected at different runs. As a result, noticeable differences between the data distributions of the training data (manually gated reference sample) and the remaining unlabeled test data (the remaining samples) are observed, and an approach such as domain-adaptation is required to remove these biases. Bead-normalization is an approach introduced to mass cytometry as a pre-processing step to diminish the effect of such variations [39]. Interestingly, application of DeepCyTOF to unnormalized and bead-normalized data did not show an advantage of using the latter for the task of automated gating. Our domain-adaptation approach allows us to effectively normalize different distributions for the (supervised learning) task of automated gating via introduction of intermediate representations of cytometry data, each consisting of instances from the reference (gated) distribution mixed with instances from a given un-gated distribution.

Flow cytometry and mass cytometry experiments provide us with multivariate data with dimensionality ranging between 10-40. Transforming the raw multivariate data to other representations may offer advantages for tasks such as automated gating or calibration. Finding good representations can be done either by manual investigation (hand crafting) or automated approaches. In recent years deep learning approaches have been shown to be suitable for learning useful representations of data in the sense that they provide new sets of features that makes subsequent learning easier. Furthermore, it has been shown that pre-training unsupervised steps such the ones we implemented in DeepCyTOF can improve the learning tasks [52], especially, when labeled training data is scarce. Cytometry experiments provide us with large datasets of unlabeled cells, which makes the unsupervised pre-training steps in the construction of a deep neural network applicable.

As cytometry analyses become widely used in research and clinical settings, automated solutions for analyzing the high dimensional datasets are urgently needed. Current practice in which samples are first subjected to manual gating are slowly substituted by automatic gating methods [53]. Major contributions to between-sample variations in cytometry experiments arise not only due to biological or medical differences but due to machine biases. Here we demonstrate that a novel machine learning approach based on deep neural networks and domain adaptation can substitute manual gating as they both produce indistinguishable results. In future work, we will demonstrate that deep learning approaches could address other challenges in analyzing cytometry data. This includes tasks such as further development of unsupervised calibration of samples, and feature extraction for classification or visualization of multiple samples.

## Acknowledgement

The authors would like to thank Catherine Blish for CyTOF reagents, Ronald Coifman, and Roy Lederman for helpful discussions and suggestions. This research was funded by NIH grant 1R01HG008383-01A1 (Y.K.).

## A Data Pre-processing

Given a blood samples A, we first perform an elementary logarithmic transformation

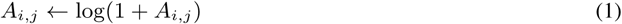

In addition, we standardize each column of A to have zero mean and and unit variance.

## Footnotes

1 Throughout this manuscript, we use the terms *sample* and *subject* as follows: a *sample* is a collection of measurements of cells, belonging to a single *subject*.

2 Our results in Section 3.3 show that this normalization procedure does not always eliminate the batch effects between different instruments, and further calibration is needed.

3 the usage of the terms “source” and “target” in this manuscript is opposite than in [37], in order to be consistent with the domain adaptation terminology.

4 the MMD values were computed using random batches of size 1000 from each of the distributions.

